# Growth patterns of theoretical bite force and jaw musculature in southern sea otters (*Enhydra lutris nereis*)

**DOI:** 10.1101/2024.08.23.609377

**Authors:** Chris J. Law

## Abstract

The transition from milk to solid food requires drastic changes in the morphology of the feeding apparatus and its performance. As durophagous mammals, southern sea otters exhibit significant ontogenetic changes in cranial and mandibular morphology to presumably enable them to feed on a variety of hard-shelled invertebrate prey. Juvenile sea otters begin feeding independently by 6 –8 months of age, but how quickly they reach sufficient maturity in biting performances remains unknown. Here, I found that theoretical bite force of southern sea otters does not reach full maturation until during the adult stage at 3.6 and 5.0 years of age in females and males, respectively. The slow maturation of biting performance can be directly attributed to the slow growth and development of the cranium and the primary jaw adductor muscle (i.e., the temporalis) and may ultimately impact the survival of newly weaned juveniles by limiting their ability to process certain hard-shelled prey. Alterative foraging behaviors such as tool use, however, may mitigate the disadvantages of delayed maturation of biting performance. In analyses of sexual dimorphism, I found that female otters reached bite force maturation earlier whereas male otters exhibit initial rapid growth in bite force–to quickly reach sufficient biting performances needed to process prey early in life–followed by a slower growth phase towards bite force maturation that coincides with sexual maturity. This biphasic growth in bite force suggests that male to male competition for resources and mates exhibits strong selection in the growth and development of skull form and function in male otters. Overall, this study demonstrates how the analysis of anatomical data can provide insight on the foraging ecologies and life histories of sea otters across ontogeny.

## Introduction

The growth and development of the vertebrate feeding apparatus often correspond to major ontogenetic shifts in diet and resource use (e.g., Herrel and O’Reilly 2006; Baliga and Mehta 2014; Gignac and Erickson 2014; Santana and Miller 2016; Higgins et al. 2018; Chatar et al. 2024). In mammals, weaning is a major ontogenetic dietary shift, as juveniles must transition from a liquid food source (i.e., milk) during the nursing stage to a diet consisting of solid foods that must be obtained and processed for survival. Unsurprisingly, the mammalian feeding apparatus undergoes tremendous ontogenetic changes in skull morphology and musculature to accommodate these dietary shifts. Ontogenetic changes in skull size (growth) and skull shape (development) facilitate the enlargement of jaw muscles and their attachment areas, increase in osteological robustness, and stronger bite force (Tanner et al. 2010; La Croix et al. 2011b; Segura 2015; Forbes Harper et al. 2017; Law et al. 2017; Law 2020; Stanchak et al. 2023). Furthermore, selection is expected to favor individuals that exhibit fast rates and/or short durations of growth and development, and thus more quickly become efficient foragers on adult diets (Herrel and Gibb 2006; Gignac and Santana 2016). The need for quick transitions to adult morphologies and performance may be particularly important for species with highly specialized diets such as carnivory or durophagy, a diet that consists of hard or tough material including bamboo, bones, exoskeletons, or shells. In carnivoran mammals, the growth and development of the carnivoran skull have been investigated by an increasing number of researchers. These ontogenetic changes (e.g. the flattening of the dorsal cranial profile, development of a more pronounced sagittal crest, broadening of the zygomatic arches, and increase in size) all contribute to stronger bite performance (Tanner et al. 2010; La Croix et al. 2011a; Segura and Prevosti 2012; Tarnawski et al. 2014; Forbes Harper et al. 2017; Chatar et al. 2024). Although adults of durophagous species exhibit strong biting performances to specialize on hard material (Binder and Van Valkenburgh 2000; Christiansen and Wroe 2007; Figueirido et al. 2014; Tseng et al. 2017), these traits are often not fully developed in the skulls of juveniles and may constrain their ability to break into their hard, tough food items. Thus, how quickly durophagous juveniles reach sufficient maturity in biting performances requires further investigation.

Southern sea otters (*Enhydra lutris nereis*) are a great model species to examine the growth and development of biting performance and the underlying jaw musculature. This durophagous marine mustelid feeds on a variety of hard-shelled invertebrates such as clams, crabs, mussels, snails, and urchins (Riedman and Estes 1990). As one of the smallest marine mammals, sea otters exhibit the highest known mass-specific metabolic rates of marine mammals (Morrison et al. 1974; Costa and Kooyman 1984; Yeates et al. 2007) and consequently must spend 20–50% of the day foraging to consume 20–30% of their body mass in food per day (Morrison et al. 1974; Costa and Kooyman 1982; Estes et al. 1986; Ralls and Siniff 1990; Thometz et al. 2016). Thus, as adults, sea otters exhibit several cranial adaptations that facilitate durophagy and enable them to feed effectively including short, blunt skulls (Riley 1985); taller and wider mandibular rami (Timm-Davis et al. 2015); and bunodont dentition with fracture-resistant dental enamel (Constantino et al. 2011; Ziscovici et al. 2014). Southern sea otters exhibit dramatic changes in cranial and mandibular shape and size over ontogeny to reach these adult morphologies including the widening of the zygomatic arches and the mastoid processes, dorsoventral compression of the cranium, shortening of the jaw length, and anterodorsal expansion of the coronoid process (Law et al. 2017). These ontogenetic changes enable southern sea otters to exhibit positive allometric increases in theoretical bite force and underlying jaw adductor musculature with respect to body size and skull size (Law et al. 2016b). Southern sea otters obtain most of their nourishment from nursing during the first few weeks of life (Payne & Jameson, 1984). However, by 6 weeks of age, pups begin soliciting small pieces of food from their mothers, and by 8.9 weeks of age, pups begin to catch and consume some of their own prey (Staedler 2011). Pups can break open some hard-shelled prey items by 14–20 weeks of age and begin to display proto-tool use behaviors (Staedler 2011). Pups are weaned and completely independent by 6–8 months of age (Payne and Jameson 1984; Jameson and Johnson 1993).

Although significant ontogenetic changes in skull morphology occur during these first 6–8 months, the full growth and development of the skull does not occur until well after weaning (Law et al. 2017). The mandible does not reach adult maturity until the subadult stage at 1–3 years old, and most aspects of cranial maturity do not occur until after mature body size is obtained during the adult stage at 4–5.5 years old (Law et al. 2017). Law et al. (2017) postulated that the slower maturation of the skull may hamper the ability of newly weaned juveniles to successfully process hard-shelled prey by constraining jaw adductor muscle size, and thus biting performance. Although positive allometry in bite force suggests fast growth to overcome the smaller, less developed morphology of the juvenile stage (Law et al. 2016), the rate and duration of growth and development of biting performance remains to be quantified.

In this study, I quantified the age at which juvenile southern sea otters reach maturity in biting performance. To accomplish this goal, I conducted dissection-based estimations of theoretical bite forces (Davis et al. 2010; Santana et al. 2010) across an ontogenetic series of southern sea otters and generated a growth curve model of theoretical bite force. I also generated growth curves for the underlying jaw adductor musculature and lever mechanics known to contribute to bite force generation in mammals. Lastly, because male sea otters attain maturation of skull morphologies earlier than female otters (Law et al. 2017), I tested whether male otters also exhibit faster maturation of theoretical bite force and underlying jaw musculature.

## Methods

### Specimens and gross dissections

Osteological and muscular data were gathered from 74 southern sea otter specimens. 55 of these specimens were obtained from Law et al. (2016b) and 19 specimens were added in this study. All 74 specimens originated from naturally deceased southern sea otters obtained from the California Department of Fish and Wildlife (CDFW) Marine Wildlife Veterinary Care and Research Center between October 2013 and June 2017. The age of each specimen was estimated by CDFW using a suite of morphological characteristics including total body length, tooth wear, and closure of cranial sutures (Nicholson et al. 2020). Ages of studied specimens ranged from 1 day to 11 years (Fig. 1). Following CDFW, I also classified each individual into one of five age classes based on their estimated age: pups (0–6 months), immatures (6–12 months), subadults (1–3.5 years), adults (3.5–10 years), and aged adults (>10 years). All specimens stranded along the central California coast, within and throughout the current southern sea otter range from Pigeon Point in the north to Gaviota in the south. All specimens were deposited at the California Academy of Sciences. Overall, my dataset consisted of 35 female otters (nine pups, six immatures, eight subadults, 10 adults, and two aged adults) and 39 male otters (seven pups, six immatures, seven subadults, 15 adults, and four aged adults).

**Fig. 1.**
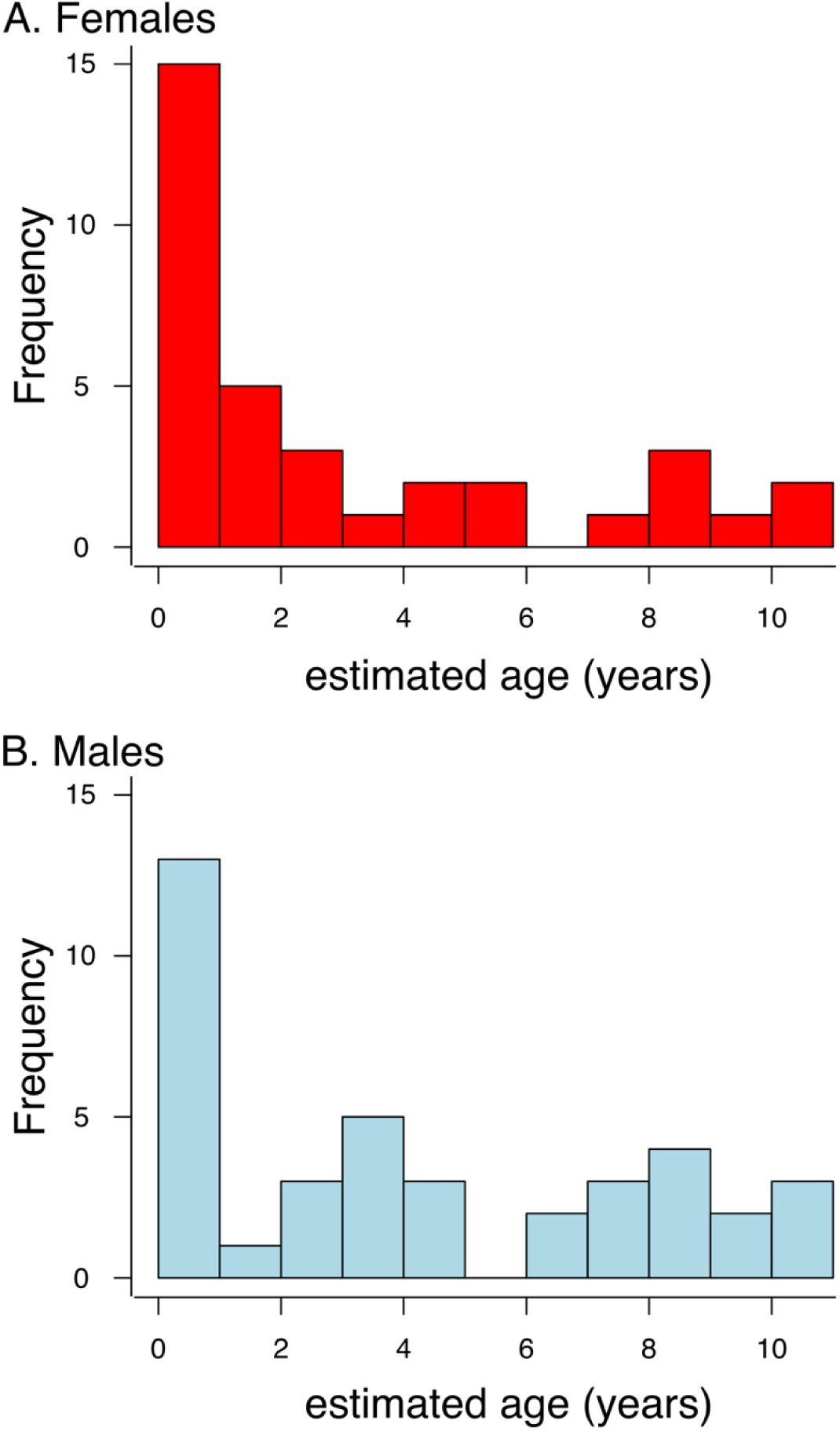
Frequency plots of sea otter specimens by estimated age (years).

I dissected the major jaw adductor muscles (superficial temporalis, deep temporalis, superficial masseter, deep masseter, and zygomaticomandibularis). While there was distinct separation between the superficial and deep temporalis, there was inadequate separation between the superficial masseter, deep masseter, and zygomaticomandibularis muscles (Scapino 1968; Law et al. 2016b). I did not dissect the medial and lateral pterygoids because these muscles are positioned deep along the medial side of the mandible and cannot be excised intact fully.

Muscles were removed from the left side of the skull, blotted dry, and weighed to the nearest 0.1 g using a digital scale. I measured the length and pennation angle of muscle fibers by digesting and separating the muscles in a solution of 15% aqueous nitric acid for 3–7 d depending on muscle size (Biewener and Full 1992). Muscle fiber lengths were measured to the nearest 0.01 mm using a digital caliper. Skulls were subsequently cleaned by a dermestid beetle colony at the California Academy of Sciences or with a maceration tank at CDFW. I then photographed and digitally measured the condylobasal length (i.e., distance from the anteriormost point on the premaxillae to the plane of the posterior surface of the occipital condyles), zygomatic breadth (i.e., greatest distance across the zygomatic arches), and cranial height (i.e., distance perpendicular to the palate plane from the lateralmost point of the mastoid process to the point of the sagittal crest directly superior to the mastoid process) of each cranium using ImageJ (Schneider et al. 2012). I then calculated the geometric mean of these three measurements as a proxy for overall skull size of each specimen. The geometric mean was derived from the Nth root of the product of N (N = 3) linear measurements.

### Estimating theoretical bite force

I calculated a theoretical bite force for each specimen by modeling the mandible as a static third-class lever (Fig. 2). In this model, an axis passing through the temporomandibular joints (TMJs) serves as a fulcrum and muscle forces generated by contractions of the temporalis and masseter adductor muscles create rotation of the lower jaw about this fulcrum (Davis et al. 2010). Under static lever equilibrium, the force of biting balances the rotation of the lower jaw created by these muscle forces. To simplify the bite force model, I treated the superficial and deep temporalis subdivisions of the temporalis as one muscle (i.e., the temporalis) and the superficial masseter, deep masseter, and zygomaticomandibularis as another muscle (i.e., the masseter). I first calculated the physiological cross sectional area (PCSA) for the temporalis and masseter muscles based on their masses (m), mean fiber lengths (f), fiber pennation angles (θ), and muscle density as a constant (ρ = 1.06 g/cm (Mendez and Keys 1960)), (Sacks et al. 1982):

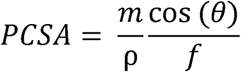

**Fig. 2.**
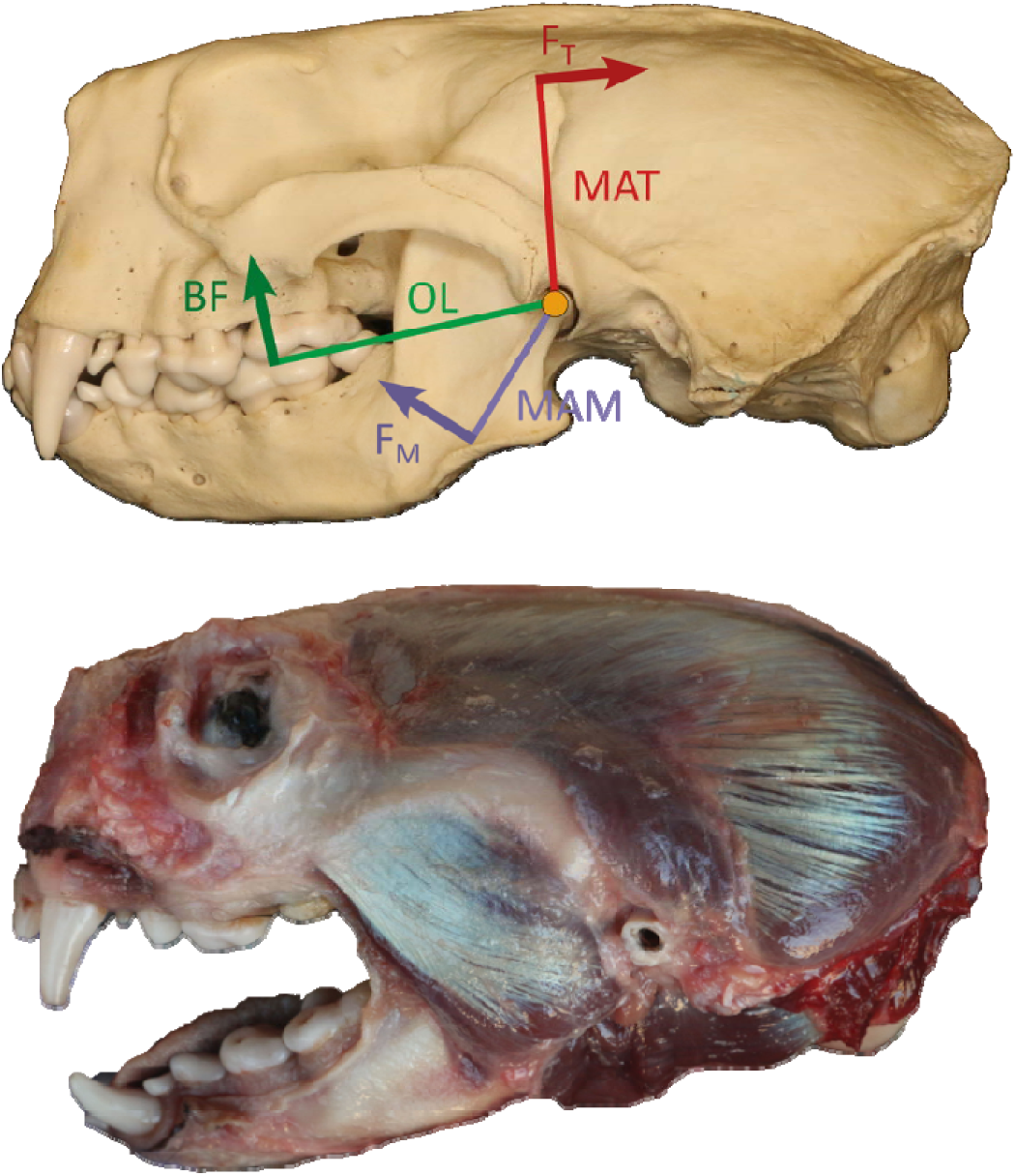
Diagram of the bite force model. Forces of the temporalis and masseter muscles (F_T_ and F_M_) are applied at given distances (MAT and MAM, respectively) from the TMJ (orange circle), creating rotation of the lower jaw, and bite force at a given distance (OL) balances these muscle forces: BF = 2[(F_T_ × MAT + F_M_ × MAM)/OL].

Pennation angle remains difficult to measure, particularly in muscle complexes that consist of multiple muscles with different lines of action. Thus, as part of my two-muscle bite force model simplification, I used a pennation angle of 0° in estimations of both muscle PCSAs.

I then estimated muscle forces of the temporalis and masseter by multiplying the PCSA with a muscle stress value of 30 N/cm^2^ (Davis et al. 2010). I modeled each muscle force as a single force vector (F_T_ and F_M_) perpendicular to the muscle in-levers and applied them to the insertion points of the temporalis (the top of the coronoid process of the mandible) and the masseter (the mid-point between the anteriormost edge of the masseteric fossa and the angular process of the mandible) (Fig. 2). The temporalis in-lever (MAT) is the in-lever length measured from the insertion point of the temporalis to the TMJ, and the masseter in-lever (MAM) is the in-lever length from the insertion point of the masseter to the TMJ. Estimations of muscle forces assumed that all jaw muscles were maximally activated.

I calculated theoretical bite forces in between the first and second molars of the lower jaw because the molars are used to crush items before consumption (Riedman and Estes 1990).

Maximum theoretical bite force (BF) was estimated by adding the moment (product of the force vector and in-lever length) of each jaw adductor muscle, dividing by the out-lever,and multiplying by 2 to account for bilateral biting:

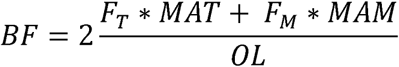

where F_T_ and F_M_ are the force vectors of the temporalis and masseter, respectively; MAT is the temporalis in-lever length; MAM is the masseter in-lever length; and OL is the out-lever measured from the bite point and the TMJ (Fig. 2). All in-lever and out-lever measurements were taken from photographs of the lateral view of the mandible in ImageJ v. 1.48 (Schneider et al. 2012). Because all muscles were dissected and measured at occlusion in all specimens, muscle force vectors were also estimated based on gape angles at occlusion (i.e., 0°). Lastly, I calculated the mechanical advantage (MA) of jaw closing based on the temporalis (MAT/OL) and masseter (MAM/OL) muscles. MA is the ratio of the length of the in-lever (MAT or MAM) divided by the length of the out-lever (OL). A relatively higher mechanical advantage suggests a jaw more optimized for stronger bite force.

### Statistical Analyses

I determined the rate and the age at which skull size, theoretical bite force, underlying muscle parameters (i.e., PCSA, mass, and fiber length), and mechanical advantage of jaw closing reached maturity using Brody growth models:

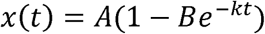

where *x(t)* is the trait of interest at time *t*, *A* is the asymptotic maturity, *k* is the exponential decay of the growth rate and *B* is the scaling constant signified by the ratio between the initial growth rate at t = 0 and *k* (Brody 1945). Thus, the initial growth rate (*L*) can be calculated as the product between *B* and *k*. I used Brody models in this study to keep the parameters consistent with previous work on sea otter skull growth that found that Brody models were the best fitting models compared to a suite of other growth models (Law et al. 2017). I fit Brody models to each trait separately by sex because southern sea otters exhibit sexual dimorphism in the skull (Law et al. 2016a, 2017). I then determined whether the parameter estimates significantly differed between the sexes by comparing the mean values of one sex with the 95% confidence intervals of the mean with the other sex. I also reported the age of maturity of each trait as the estimated age at which the trait reached the lower limit of the 95% confidence interval for its asymptotic value (Zelditch et al. 2003). All growth models were fit using the nls function in R 4.3.1 (R Core Team 2023).

## Results

Asymptotic values of theoretical bite force significantly differed between male and female southern sea otters; adult males exhibited 142% greater theoretical bite forces (BF_♀_ = 437.7 N; BF_♂_ = 620.7 N) than adult females (Table 1; Fig. 3). Although females displayed faster initial growth rates, males displayed slower rates of growth decay to reach their theoretical bite force, leading to later maturation (i.e., 95% of the asymptotic values) in theoretical bite force (5.0 years) compared to females (3.6 years). However, males reached the asymptotic value of theoretical bite force of females in 1.8 years.

**Fig. 3.**
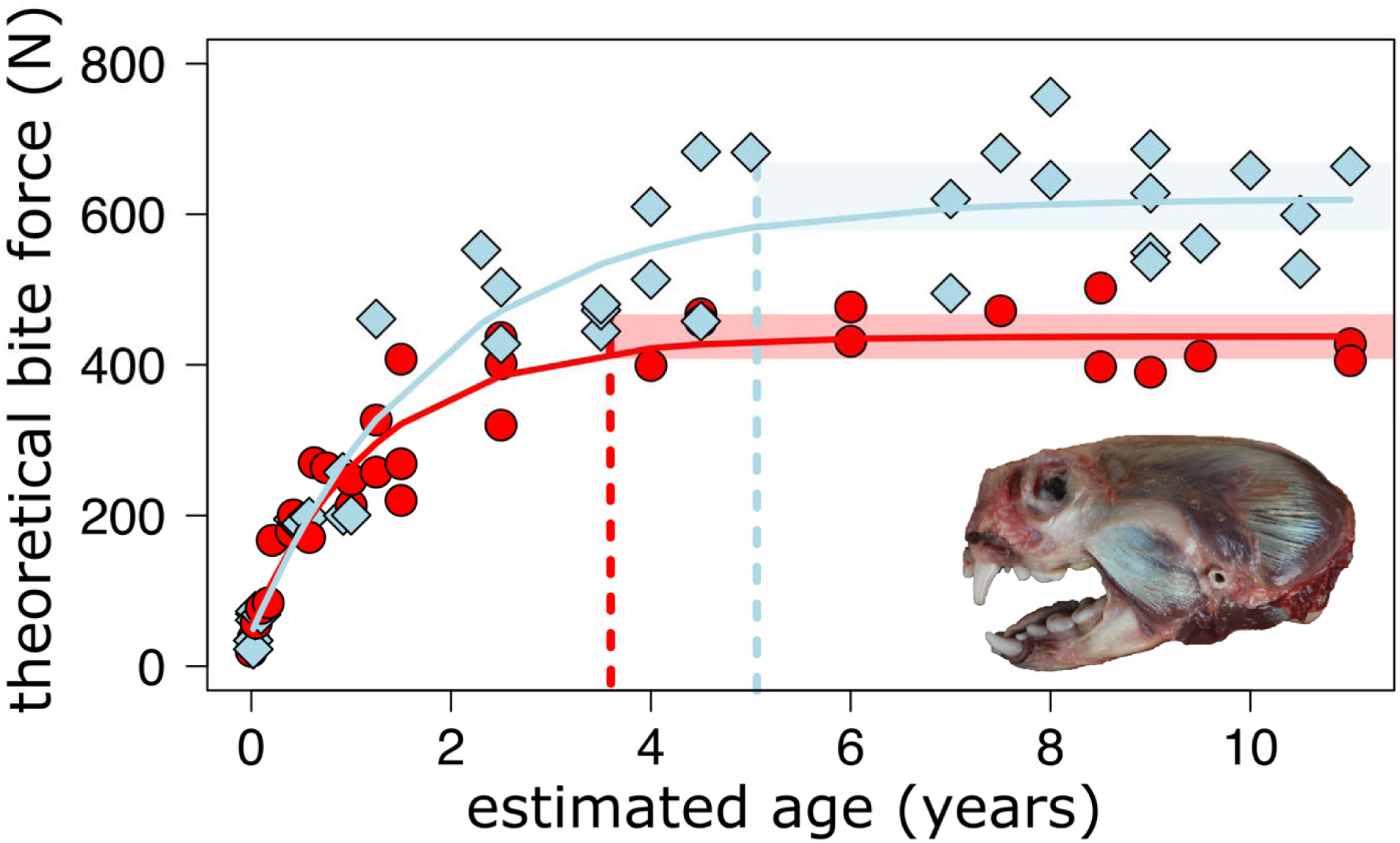
Growth curve for theoretical bite force. Red circles and growth curves represent female otters and blue diamonds and growth curves represent male otters. Red and blue dashed lines indicate the times at which traits reach 95% of their asymptotic values for females and males, respectively.

**Table 1.**
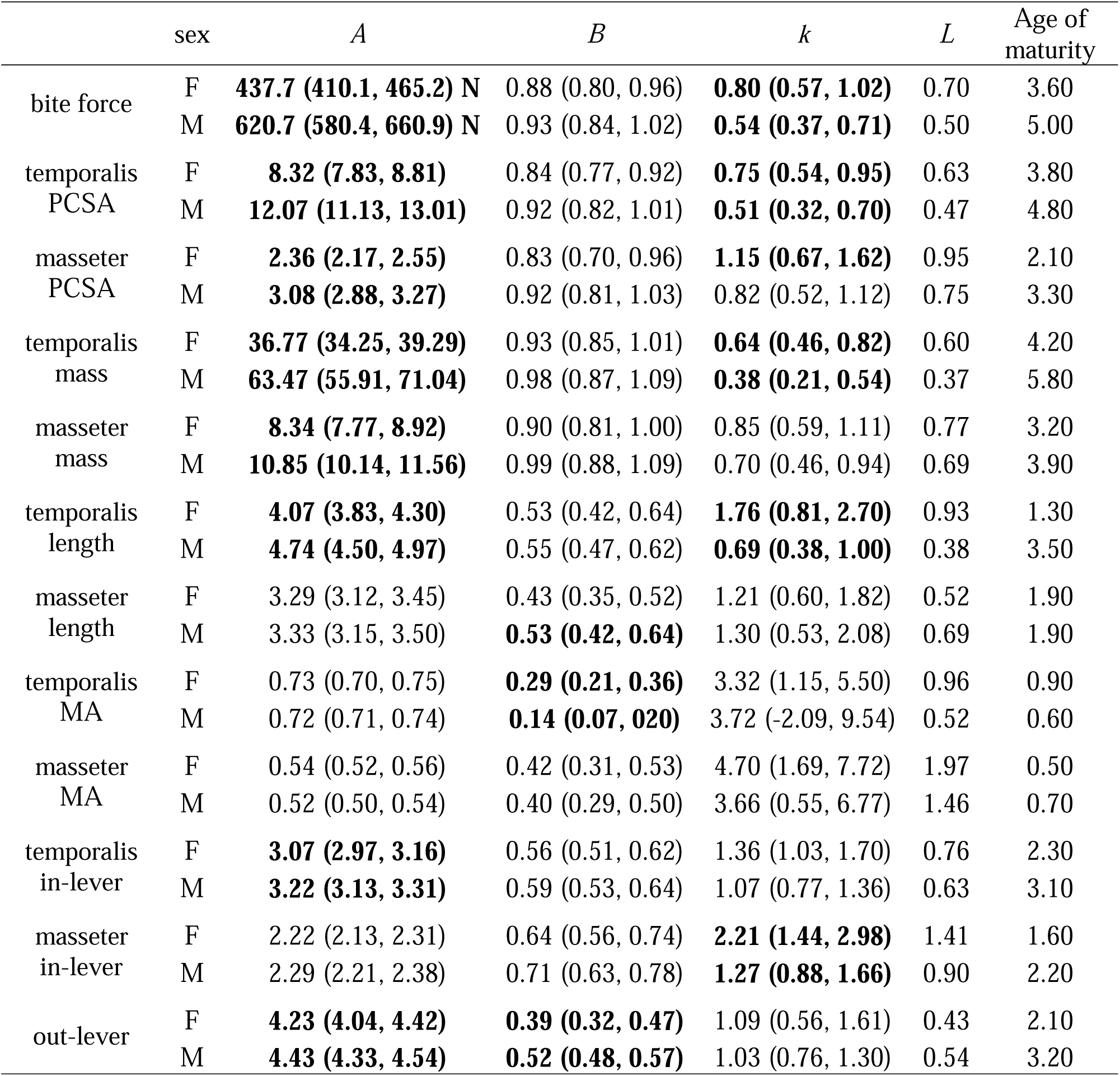
Parameter estimates of growth curves generated with a Brody model. *A* is the asymptotic maturity, *k* is the exponential decay of the growth rate, *B* is the scaling constant signified by the ratio between the initial growth rate at t = 0 and *k*, and *L* is the initial growth rate calculated as the product between *B* and *k*. 95% confidence intervals are given in parentheses. Bold values indicate significant differences between male and female parameters. Age of maturity indicates the ages (in years) at which element reach 95% of their respective asymptotic values.

|  | sex | *A* | *B* | *k* | *L* | Age of maturity |
| --- | --- | --- | --- | --- | --- | --- |
| bite force | F | **437.7 (410.1, 465.2) N** | 0.88 (0.80, 0.96) | **0.80 (0.57, 1.02)** | 0.70 | 3.60 |
|  | M | **620.7 (580.4, 660.9) N** | 0.93 (0.84, 1.02) | **0.54 (0.37, 0.71)** | 0.50 | 5.00 |
| temporalis PCSA | F | **8.32 (7.83, 8.81)** | 0.84 (0.77, 0.92) | **0.75 (0.54, 0.95)** | 0.63 | 3.80 |
|  | M | **12.07 (11.13, 13.01)** | 0.92 (0.82, 1.01) | **0.51 (0.32, 0.70)** | 0.47 | 4.80 |
| masseter PCSA | F | **2.36 (2.17, 2.55)** | 0.83 (0.70, 0.96) | **1.15 (0.67, 1.62)** | 0.95 | 2.10 |
|  | M | **3.08 (2.88, 3.27)** | 0.92 (0.81, 1.03) | 0.82 (0.52, 1.12) | 0.75 | 3.30 |
| temporalis mass | F | **36.77 (34.25, 39.29)** | 0.93 (0.85, 1.01) | **0.64 (0.46, 0.82)** | 0.60 | 4.20 |
|  | M | **63.47 (55.91, 71.04)** | 0.98 (0.87, 1.09) | **0.38 (0.21, 0.54)** | 0.37 | 5.80 |
| masseter mass | F | **8.34 (7.77, 8.92)** | 0.90 (0.81, 1.00) | 0.85 (0.59, 1.11) | 0.77 | 3.20 |
|  | M | **10.85 (10.14, 11.56)** | 0.99 (0.88, 1.09) | 0.70 (0.46, 0.94) | 0.69 | 3.90 |
| temporalis length | F | **4.07 (3.83, 4.30)** | 0.53 (0.42, 0.64) | **1.76 (0.81, 2.70)** | 0.93 | 1.30 |
|  | M | **4.74 (4.50, 4.97)** | 0.55 (0.47, 0.62) | **0.69 (0.38, 1.00)** | 0.38 | 3.50 |
| masseter length | F | 3.29 (3.12, 3.45) | 0.43 (0.35, 0.52) | 1.21 (0.60, 1.82) | 0.52 | 1.90 |
|  | M | 3.33 (3.15, 3.50) | **0.53 (0.42, 0.64)** | 1.30 (0.53, 2.08) | 0.69 | 1.90 |
| temporalis MA | F | 0.73 (0.70, 0.75) | **0.29 (0.21, 0.36)** | 3.32 (1.15, 5.50) | 0.96 | 0.90 |
|  | M | 0.72 (0.71, 0.74) | **0.14 (0.07, 0.20)** | 3.72 (-2.09, 9.54) | 0.52 | 0.60 |
| masseter MA | F | 0.54 (0.52, 0.56) | 0.42 (0.31, 0.53) | 4.70 (1.69, 7.72) | 1.97 | 0.50 |
|  | M | 0.52 (0.50, 0.54) | 0.40 (0.29, 0.50) | 3.66 (0.55, 6.77) | 1.46 | 0.70 |
| temporalis in-lever | F | **3.07 (2.97, 3.16)** | 0.56 (0.51, 0.62) | 1.36 (1.03, 1.70) | 0.76 | 2.30 |
|  | M | **3.22 (3.13, 3.31)** | 0.59 (0.53, 0.64) | 1.07 (0.77, 1.36) | 0.63 | 3.10 |
| masseter in-lever | F | 2.22 (2.13, 2.31) | 0.64 (0.56, 0.74) | **2.21 (1.44, 2.98)** | 1.41 | 1.60 |
|  | M | 2.29 (2.21, 2.38) | 0.71 (0.63, 0.78) | **1.27 (0.88, 1.66)** | 0.90 | 2.20 |
| out-lever | F | **4.23 (4.04, 4.42)** | **0.39 (0.32, 0.47)** | 1.09 (0.56, 1.61) | 0.43 | 2.10 |
|  | M | **4.43 (4.33, 4.54)** | **0.52 (0.48, 0.57)** | 1.03 (0.76, 1.30) | 0.54 | 3.20 |

Asymptotic values of the temporalis PCSA, mass, and fiber lengths also significantly differed between males and females (Table 1; Fig. 4). Adult male otters exhibited greater temporalis PCSA, larger temporalis mass, and longer temporalis fiber lengths but displayed slower initial growth rates (significantly lower decay rates with similar scaling constants) and thus later maturation in these traits compared to adult female otters (Table 1). Adult male otters also exhibited greater masseter PCSA and larger masseter mass than adult female otters; however, neither scaling constants nor rates of growth decay in these masseter traits do not significantly differ between the sexes, suggesting that increased growth duration rather than slower growth rate contributed to the larger PCSA and mass found in males. In contrast, asymptotic values of masseter fiber lengths did not significantly differ between the sexes.

**Fig. 4.**
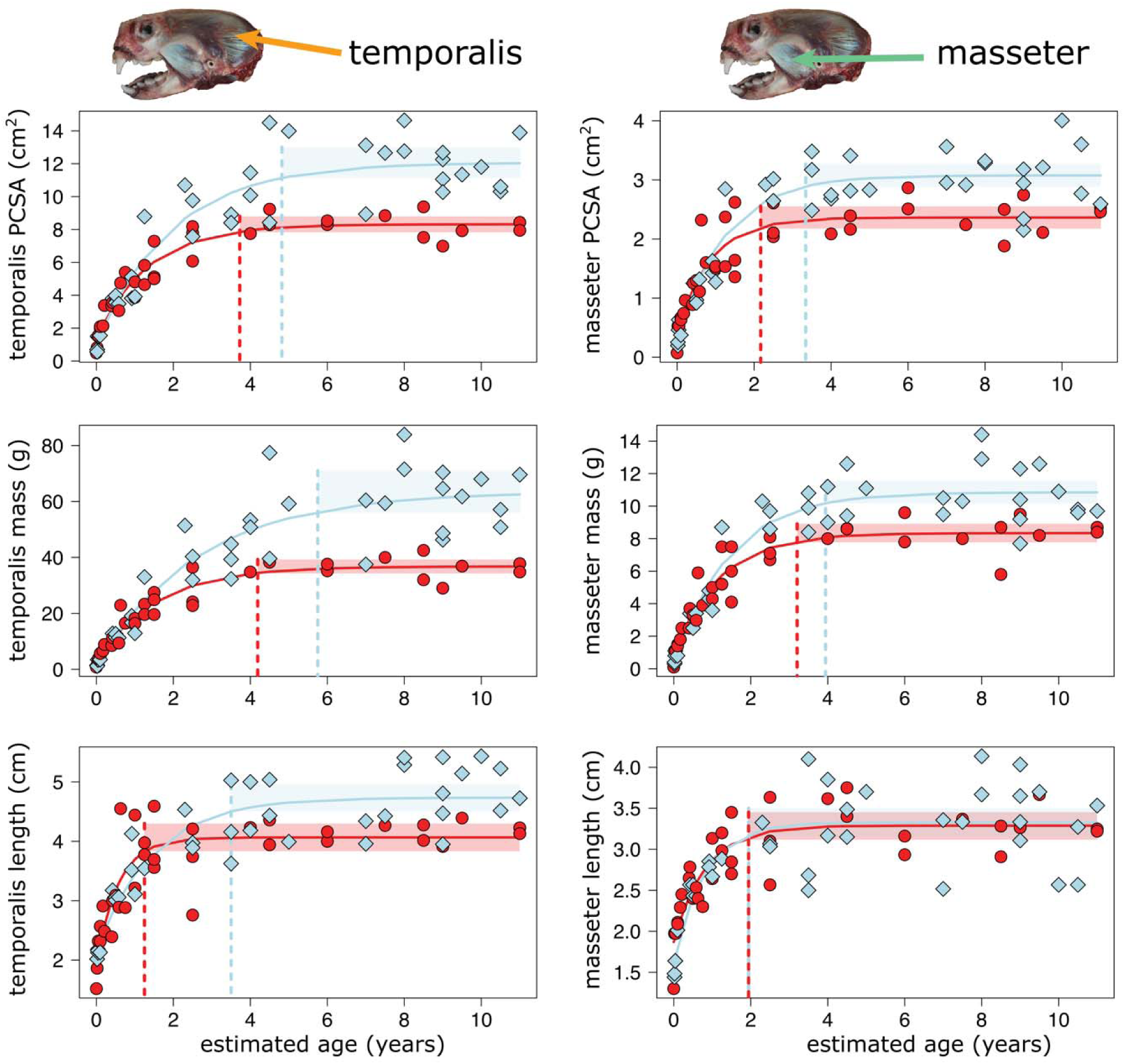
Growth curves for PCSA, mass, and fiber length of the temporalis and masseter muscles. Red circles and growth curves represent female otters, and blue diamonds and growth curves represent male otters. Red and blue dashed lines indicate the times at which traits reach 95% of their asymptotic values for females and males, respectively.

Asymptotic values and rates of growth decay of the mechanical advantage of jaw closing for either the temporalis or masseter did not differ between males and females (Table 1; Fig. 5). Both female and male otters reached mature mechanical advantage during weaning (6–12 months of age). The 95% confidence interval of the *k* parameter in the model for temporalis mechanical advantage in males includes 0, suggesting that the Brody model may be a poor fit to this data. Asymptotic values of temporalis in-lever (MAT) and out-lever to the molars (O) differed between the sexes; adult male otters exhibited significantly longer MAT and OL than females (Table 1; Fig. 5). Interestingly, the temporalis in-lever of males reached maturation earlier than the out-lever of males by over a year, which may contribute to the poor fit of the *k* parameter. In contrast, rates of growth decay of masseter in-lever (MAM) differed between the sexes but not their asymptotic values (Table 1; Fig. 5).

**Fig. 5.**
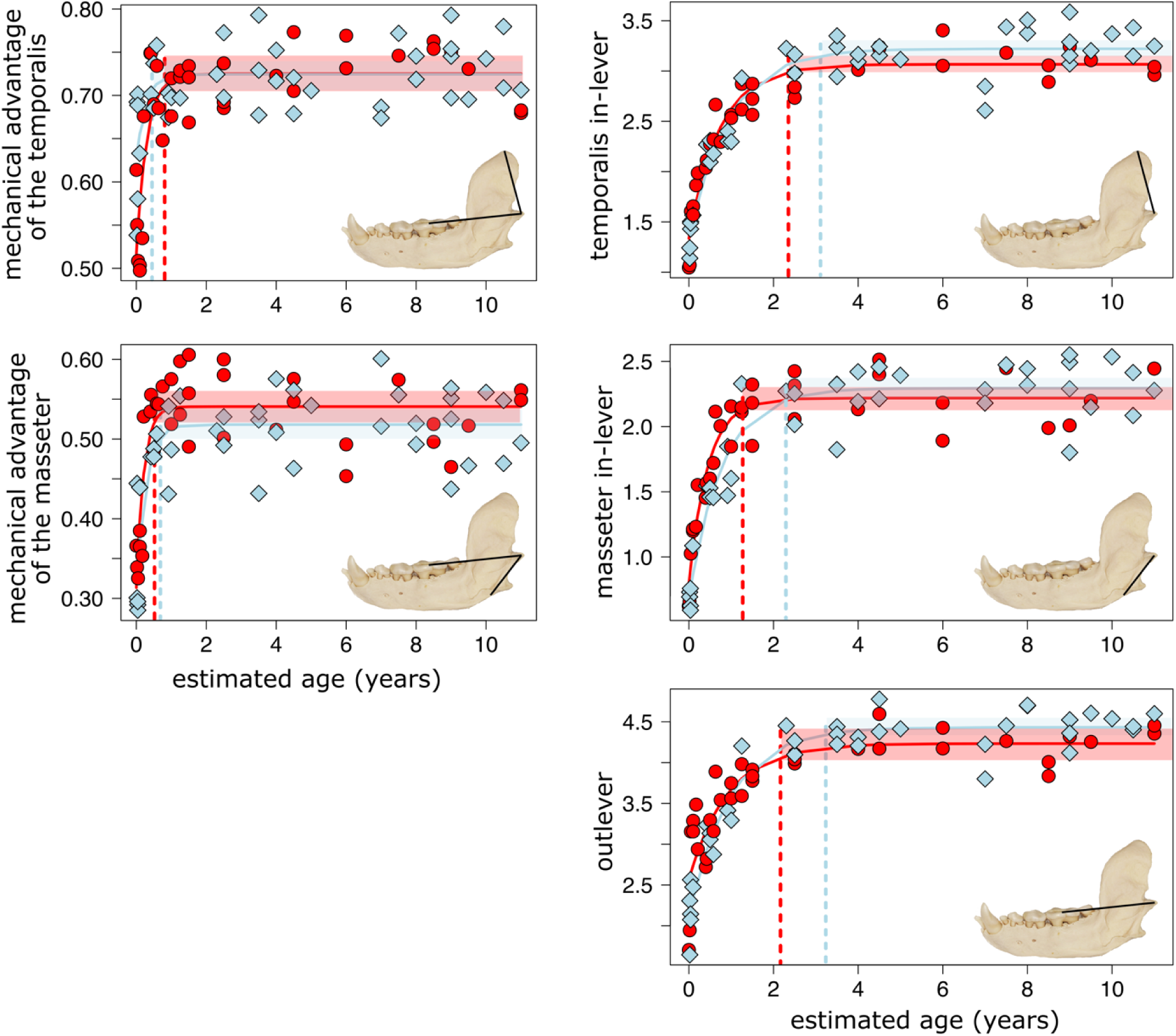
Growth curves for mechanical advantage of jaw closure from the temporalis and masseter muscles. Red circles and growth curves represent female otters and blue diamonds and growth curves represent male otters. Red and blue dashed lines indicate the times at which traits reach 95% of their asymptotic values for females and males, respectively.

## Discussion

The transition from milk to a diet consisting of hard-shelled invertebrate prey presumably requires equally drastic changes in the morphology of the feeding apparatus and biting performance. In this study, I found that bite force does not reach full maturity until the adult stage (3.6 years old in females and 5.0 years old in males). The slow maturation of biting performance may limit newly weaned juveniles from processing certain hard-shelled prey and may therefore impact their survival. The first year post-weaning age class often shows depressed survivorship (Monson et al. 2000; Tinker et al. 2006), as newly weaned otters exhibit the highest daily energy demands of all juvenile age classes (Thometz et al. 2014) but do not have the benefit of obtaining food from their mothers and must compete with conspecific adults for the same food resources (Riedman and Estes 1990) in already resource-poor environments (Tinker et al. 2008; Layman et al. 2015; Tinker and Hatfield 2015). Together, these results suggest that newly weaned southern sea otters are at a disadvantage in foraging until the early stages of adulthood when they reach adult skull morphologies and biting performances. Below, I discuss how alternative feeding behaviors may help mitigate delayed maturation of biting performance and how craniomandibular asynchronization and sexual dimorphism influence bite performance growth. Lastly, I compare how sea otter bite force growth compares to other carnivorans and discuss caveats to my bite force model.

### Tool use may mitigate delayed maturation of biting performance

Alternative feeding behaviors can potentially provide pathways to foraging success even with suboptimal morphologies or performances such as a less developed feeding apparatus. One such feeding behavior found in southern sea otters is the use of rocks, shells, or other hard objects as hammers or anvils to crack open their hard-shelled prey (Riedman and Estes 1990).

The use of tools varies greatly among individual otters and is associated with individual variation in diet specialization; for instance, adult otters specializing on snails or clams tend to use tools frequently whereas adult otters specializing on urchins, crabs, and small bivalves tend not to use tools (Fujii et al. 2015, 2017). Overall, individuals that used tools frequently processed harder, larger prey items that would be typically inaccessible with biting alone (Law et al. 2024b). These benefits may be enhanced in juvenile otters as they exhibit weaker bite forces compared to their adult conspecifics. All sea otter pups exhibit proto-tool use behaviors (such as repetitive pounding of objects on their chests) and begin using tools at the age of 14 weeks (Staedler 2011). During this stage, an average otter’s bite force is only 88 N, far lower than the amount of force it takes to break open even the least hard prey such as urchins (mean fracture force = 185 N) and crabs (mean fracture force = 247 N) (Law et al. 2016b, 2024b). Tool use behavior does not vary with the mother’s diet specialization (Staedler 2011). Thus, although urchin and crab specialists tend not to use tools as adults, juvenile otters use tools to process these prey types (Staedler 2011), suggesting that tool use may contribute to their foraging success and compensate for their weaker bite forces. Furthermore, because larger prey items contain more calories but are harder to break open (Law et al. 2024b), tool use may also enable juveniles to gain access to additional calories that would have been inaccessible with their immature bite forces. These tool use benefits may be particularly important for female otters as they used tools with greater frequencies than male otters, which enabled them to consume prey items that were 21–35% harder compared with males despite their smaller crania and weaker bite forces (Law et al. 2024b). Therefore, it is tempting to postulate that tool use may serve as an essential behavior in facilitating prey access for individuals with weak bite forces that are still growing/developing or are smaller. Nevertheless, the majority of tool use research in sea otters has focused on adults and additional work is required to further understand the importance of tool use on foraging success and survival in juvenile otters.

### Effects of craniomandibular asynchronization on biting performance growth

Asynchronized growth and development between the cranium and mandible found in southern sea otters may explain the underlying factors that are affecting the maturation rate of biting performance. Compared to the mandible, the sea otter cranium exhibits slower growth and development of its morphology (Fig. 6) (Law et al. 2017). Because the cranium serves as the underlying structure that supports the primary jaw muscle (i.e., the temporalis muscle) used to generate bite force, the slow maturation of cranial size and shape directly contributes to the slow maturation of biting performance (3.6 years of age for females and 5.0 years of age for males).

**Fig. 6.**
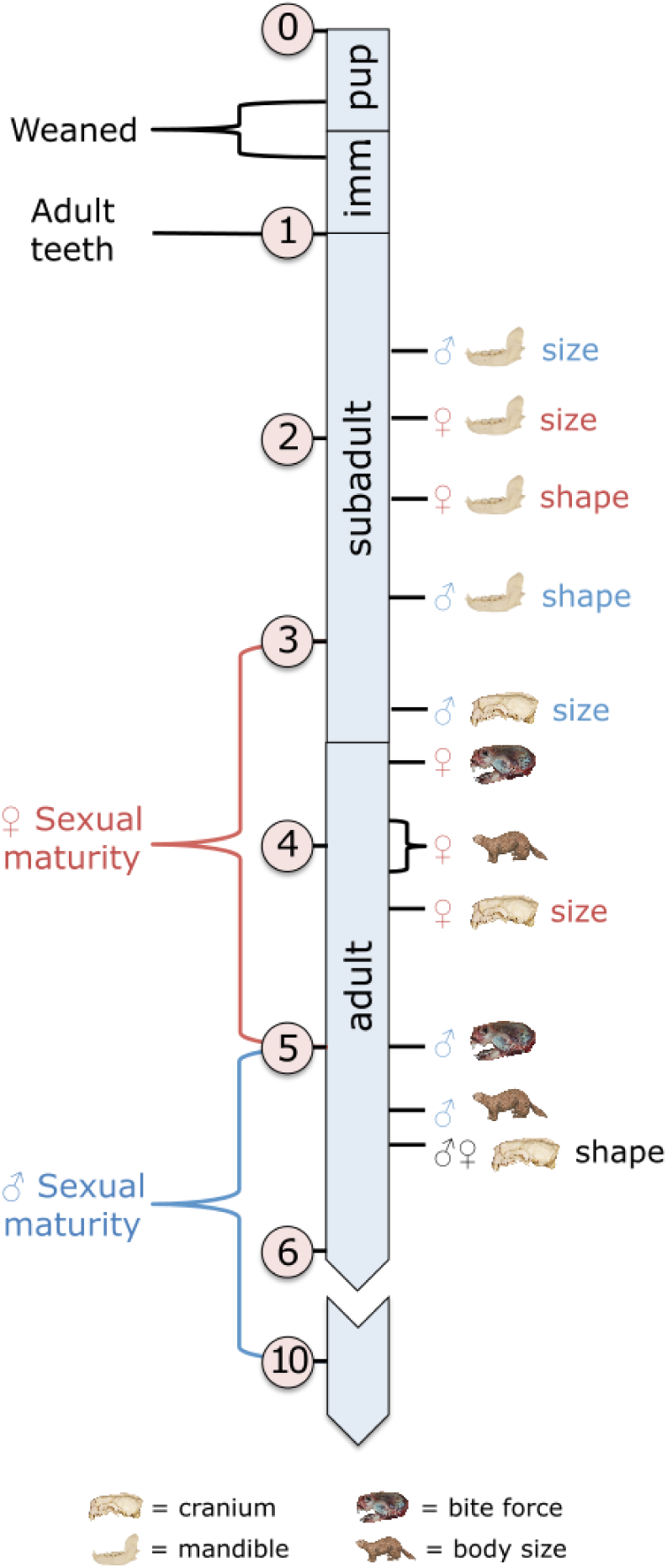
Timeline illustrating the maturation ages of southern sea otter skull morphology and theoretical bite force in relation to major life-history events. Maturation ages of cranial and mandibular size and shape were obtained from Law et al. (2017). Cranial and mandibular size and shape were calculated using geometric morphometrics. Maturation age was defined as the age at which the trait reached 95% of their asymptotic values. Estimated age (in years) are in circles. Imm = immature pup.

Specifically, broadening of the zygomatic arches, lengthening along the anteroposterior axis, and development of a more pronounced sagittal crest all provide additional attachment sites for the temporalis muscle (Scapino 1968; Turnbull 1970; Tanner et al. 2010; La Croix et al. 2011b; Law and Mehta 2019; Law 2020; Stanchak et al. 2023). Cranial size maturity is not reached until 4.3 years of age for females and 3.3 years of age for males, cranial shape maturity is not reached until 5.5 years of age for both sexes (Fig. 6; Table 1; Law et al. 2017), and correspondingly, physiological cross-sectional area of the temporalis does not reach full maturity until 3.8 years of age for females and 4.8 years of age for males (Fig. 4; Fig. 6; Table 1). In contrast, mandibular size and shape reach full maturity during the subadult class between 1.5 to 2.8 years of age, much earlier than the cranium (Fig. 6; Table 1; Law et al. 2017). Ontogenetic changes in mandibular shape of southern sea otters are associated with increasing jaw adductor muscle in-levers and the out-lever (Law et al. 2017); thus, these lever measurements along with mechanical advantage of jaw closure also reach full maturity during the subadult class (Fig. 5; Fig. 6; Table 1). The mandible also serves as the underlying structure that supports the masseter muscle.

Unsurprisingly, physiological cross-sectional area of the masseter also reaches full maturity at 2.1 years of age for females and 3.3 years of age for males, on par with the mandibular morphologies (Fig. 4; Table 1). The earlier maturation of mandibular morphology and musculature in juvenile otters may contribute to positive allometric increases in bite force early in ontogeny (Law et al. 2016b) and provide a boost in bite force during a period in which the cranium remains smaller and less developed and the temporalis has not yet reached full development. Similar patterns (i.e., earlier maturation of mechanical advantage and mandible compared to the cranium) are also found in the bone-eating spotted hyena (Tanner et al. 2010), providing evidence that adaptations to durophagy suggest fast maturation of jaw mechanics while waiting for the rest the skull and jaw musculature to reach full size and shape.

The different functional roles of the cranium and mandible may also influence the asynchrony between their growth and/or development. The cranium is the more complex structure of the two, serving as a multifunctional structure that takes part in sensory functions, respiration, and brain protection in addition to feeding, whereas the mandible is involved primarily in feeding. Thus, the cranium may require slow, prolonged growth and/or development of its morphologies to accommodate its multiple functions. Additional studies are needed to reveal the maturation ages of growth and development among the cranium, brain, and other sensory organs in sea otters and other mammals.

Although asynchrony has been quantified in only a handful of mammals, primarily carnivorans (Tanner et al. 2010; La Croix et al. 2011b; Segura et al. 2013; Segura 2015; Law et al. 2017), this phenomenon has also been uncovered at the macroevolutionary level. Decoupled evolutionary modes between the cranium and mandible are found in various mammal clades (McLean et al. 2018; Arbour et al. 2019; Michaud et al. 2020; Cassini and Toledo 2021; Law et al. 2022, 2024a) despite strong integration between the cranium and mandible (Hautier et al. 2012; Figueirido et al. 2013; McLean et al. 2018; Michaud et al. 2020; Law et al. 2024a). A possible explanation is that the cranium exhibits more structural, functional, or phylogenetic constraints on its evolution, whereas the mandible may evolve more directly in response to dietary changes. Across carnivorans, cranial shape follows clade-based evolutionary shifts, suggesting that cranial diversity is phylogenetically constrained in carnivorans. In contrast, mandibular shape evolution is linked to broad dietary regimes, suggesting greater evolutionary lability of the mandible (Figueirido et al. 2011; Law et al. 2022, 2024a). Similarly, in bats, sensory functions (i.e., echolocation and vision) and phylogenetic structure are the most influential factors shaping cranial evolution, whereas diet has a stronger influence on mandibular evolution (Arbour et al. 2019). The comparison between asynchronous growth/development at the species level and decoupled evolution at the macroevolutionary level presents itself as an exciting avenue of future investigation in the factors influencing cranial and mandibular variation.

### Effects of sexual dimorphism on biting performance growth

Like many species within the order Carnivora (Lindenfors et al. 2007; Law 2019), a combination of sexual selection and niche divergence is hypothesized to drive sexual dimorphism in sea otters in which males are bigger. Despite their larger size, male southern sea otters displayed faster initial growth and developmental rates in skull size and some cranial shape elements compared to females (Law et al. 2017). Presumably, reaching maturation of craniomandibular morphology earlier enables males to reach maximum biting performance faster, which may lead to competitive advantages in processing hard-shelled prey and in male– male agonistic interactions. However, the growth curves of theoretical bite force provide a more nuanced pattern that displayed both faster and slower growth (Fig. 3; Table 1). Within their first two years, male otters reached the asymptotic value of theoretical bite force of females (437.7 N) in just 1.8 years compared to 3.6 years for females. Because of their lower rate of growth decay, subsequent growth of biting performance slows down and takes an additional >3 years for males to reach their asymptotic value of theoretical bite force (620.7 N). Biphasic growth of biting performance facilitated by lower rate of growth decay may be advantageous for males, which must compete to establish reproductive territories, aggressively exclude other mature males, and maintain exclusive access to estrous females (Ames et al. 1983; Jameson 1989; Ralls et al. 1996; Pearson and Davis 2005). In the first phase, quickly reaching biting performances on par with female bite forces enables males to compete with conspecifics in obtaining and processing the energy required to further grow and develop. In the second phase, continual growth of biting performance is slowed, likely because muscle tissue is energetically costly to grow (Millward et al. 1976). Consistently, the mass of the temporalis muscles is the last element associated with the feeding apparatus to reach full maturation (Fig. 4; Table 1). During this period, males do not have territories and instead congregate in ‘bachelor areas’ where they take part in extensive male–male interactions ranging from playful to aggressive that may be important for the development of fighting skills and possibly even the establishment of social hierarchies (Jameson 1989; Lafferty and Tinker 2014). Because biting is the primary weapon used in territorial conflicts between males, males that attain stronger biting performances faster may rise through the hierarchical ranks and begin to venture out of bachelor groups to establish their own reproductive territories. The ability to maintain these territories has lifetime reproductive consequences as males that maintained their territories for the longest sired the most pups (Tarjan and Tinker 2016). That full maturation of theoretical bite force and the onset of sexual maturity and body size maturity occur at the same time at five years of age in males suggest biting performance has an important influence on reproductive success.

Female otters, in contrast, exhibit a faster rate of growth decay, leading to weaker bite forces compared to male otters. A possible explanation for this pattern is that selection for larger bite forces is relaxed once female otters attained sufficient bite force to process hard prey as they do not need stronger bites and larger body sizes for territorial conflicts. Rather than for growth, energetic resources may be used to maintain their extremely high daily total energetic demands associated with raising a pup (Thometz et al. 2014, 2016). Furthermore, these lower bite forces may be sufficient to process hard-shelled prey because female otters tend to use tools more frequently than male otters, providing them an alternative method to forage effectively (Law et al. 2024b).

### Comparisons of bite force growth to other carnivorans

Despite a large body of work examining the growth and development of the carnivoran skull (Tanner et al. 2010; La Croix et al. 2011a; Segura and Prevosti 2012; Tarnawski et al. 2014; Forbes Harper et al. 2017; Chatar et al. 2024), the growth of bite force and underlying masticatory muscles is rarely studied in carnivorans. The most likely explanation is that performance and muscular data are difficult to obtain in carnivorans, much less across an entire ontogenetic series ranging from day old pups to years old adults. Thus, the few studies examining performance across ontogeny have relied on osteological specimens to estimate biting performance across ontogeny such as with calculations of mechanical advantage of jaw closing or estimations of bite force using the dry skull method (Tanner et al. 2010; La Croix et al. 2011b; Segura and Prevosti 2012; Forbes Harper et al. 2017). However, as seen in previous work, mechanical advantage reaches maturation ages much faster than other metrics of biting performance and thus may be unreliable to assess performance maturation by itself (Tanner et al. 2010; La Croix et al. 2011b); this study). Similarly, bite forces estimated with the dry skull method remain to be validated with empirical data (see *Caveats to bite force model* section below). Dissection-based models may provide better insight on biting performance growth, but muscular data remains difficult to gather across a full ontogenetic series in carnivorans. For example, Law (2020) examined ontogenetic patterns of theoretical bite forces and jaw musculature in American martens and fishers; however, this study was limited to just subadult and adult age classes and was unable to generate growth curves. Lastly, there have been very few published successes in measuring empirical bite forces in carnivorans using force transducers.

Binder and Van Valkenburgh’s (2000) examination of spotted hyenas remains the only study to successfully measure empirical bite force across ontogeny in a carnivoran. In contrast to sea otters, spotted hyena bite forces continued to increase well past skull maturity; spotted hyena skulls reached maturity around 3 years of age and their bite forces reached maturity later around 5–6.5 years of age (Binder and Van Valkenburgh 2000; Tanner et al. 2010), whereas sea otter bite forces reached maturity 3.6–5 years of age (depending on sex) prior to skull maturation at 5.5 years of age (Fig. 6). Comparison of growth rates of the underlying masticatory muscles may elucidate why bite force maturation ages differ between these two durophagous species, but this data remains to be collected in hyenas. La Croix et al. (La Croix et al. 2011b) also attempted to measure bite forces in coyotes but found that the juveniles were unmotivated biters, thus highlighting the difficulties of measuring empirical bite force across ontogeny. Overall, this study emphasizes the clear need for additional anatomical work to investigate ontogenetic relationships among skull morphology, bite performance, and the underlying masticatory musculature in carnivorans.

### Caveats to bite force model

I acknowledge that my approach of using a dissection-based 2D model with single points for each muscle force vector may be an oversimplification of bite force estimation. First, my model does not account for how different degrees of gape can influence the length and orientation of moment arms, which in turn can influence how the muscle force vectors are oriented. Increased gape is typically linked to reduced bite force (Herring and Herring 1974; Dumont and Herrel 2003; Santana 2015), suggesting that my theoretical bite force estimations may be at the upper limits. Second, the use of 3D methods may provide better estimations of bite force than 2D models. In an analysis across various bat species, Davis et al. (2010) found that 3D distributed traction models that distribute forces over each muscle’s respective attachment region on the skull is the best predictor of empirical bite force. Interestingly, the authors also found that 2D models using estimated muscle PCSAs (i.e., Thomason’s dry skull method (Thomason 1991)) do a pretty good job of estimating empirical bite forces in bats, whereas theoretical bite forces estimated using dissection-based 2D models that incorporate measured muscle PCSAs (as done in this current study) tend to be the worst of the three models as it underestimated empirical bite forces (Davis et al. 2010). These differences in bite force models may be restricted to bats, as Law and Mehta (2019) found an opposite pattern in which bite forces estimated from Thomason’s dry skull method underestimated bite forces derived from 2D dissection-based models. However, the dry skull method remains valuable in estimating muscle mass and PCSA when gross muscle is unavailable, at least for carnivorans. Dickinson et al. ((Dickinson et al. 2021)) found that the dry skull method is the best technique to estimate muscle mass and PCSA across carnivorans compared to other 2D or 3D approaches that estimate muscle attachment areas. Therefore, a 3D distributed traction model may not vastly outperform the 2D dissection-based model or Thomason’s dry skull method in sea otters; nevertheless, how these three models compare in sea otters requires additional investigation. Overall, the validity of bite force models in sea otters remains tricky to elucidate without empirical bite force measurements from wild southern sea otters. Unfortunately, direct measurements of bite forces are difficult to obtain from mammals outside of small rodents and bats (Herrel et al. 2008; Santana et al. 2010; Becerra et al. 2014), and especially from mammals protected under the US Marine Mammal Protection Act and the U.S. Endangered Species Act, as the author of this paper discovered after five years of trying.

## Conclusion

The ability to feed is paramount to survival; therefore, it is critical to understand the growth and development of the feeding apparatus and its impact on the maturation of biting performance. In this study, I found that southern sea otters exhibit slow maturation of theoretical bite force due to slow growth and development of the cranium and the temporalis muscle.

Although slow bite force maturation may limit newly weaned juveniles from processing certain hard-shelled prey and may thus impact their survival, tool using behavior may help mitigate these limitations. In addition, faster growth and development of the mandible and mechanical advantage may also provide a boost in biting performance during a period in which the cranium and temporalis has not yet reached full development. Furthermore, I found that bite force maturation ages differed between the sexes: female otters reached bite force maturation earlier whereas male otters exhibit biphasic growth in bite force in which they quickly reach sufficient bite forces needed to process prey early in life followed by a slower growth phase towards bite force maturation that coincides with sexual maturity. These patterns suggest that male to male competition for resources and mates exhibits a strong selective force in the growth and development of skull form and function in male otters. Overall, this study demonstrates how the analysis of anatomical data can provide insight on the foraging ecologies of sea otters across ontogeny. Future work integrating the functional morphology of the feeding apparatus, biomechanics of tool using behavior, and changes in prey characteristics will elucidate the processes that improve feeding performances through ontogeny.

## Acknowledgements

I am grateful to the staff of the California Department of Fish & Wildlife (Colleen Young, Erin Dodd, and Francesca Batac) and California Academy of Sciences (Moe Flannery and Sue Pemberton) for access to specimens; Lilian Carswell (US Fish and Wildlife Service) for permitting logistics; and Tim Tinker (US Geological Survey) and Rita Mehta (University of California Santa Cruz) for helpful discussions on sea otter life history. Funding was provided partly by a Grant-in-Aid of Research from the American Society of Mammalogists, a Lerner-Gray Fund through the American Museum of Natural History, a Rebecca and Steve Sooy Graduate Research Fellowship for Marine Mammals, and the National Science Foundation (NSF DBI-2128146).

